# Root-specific reduction of cytokinin perception enhances shoot growth with an alteration of *trans*-zeatin distribution in *Arabidopsis thaliana*

**DOI:** 10.1101/2021.09.02.458681

**Authors:** Kota Monden, Mikiko Kojima, Yumiko Takebayashi, Takamasa Suzuki, Tsuyoshi Nakagawa, Hitoshi Sakakibara, Takushi Hachiya

**Affiliations:** Department of Molecular and Functional Genomics, Interdisciplinary Center for Science Research, Shimane University, Matsue 690-8504, Japan; RIKEN Center for Sustainable Resource Science, 1-7-22, Suehiro, Tsurumi, Yokohama 230-0045, Japan; College of Bioscience and Biotechnology, Chubu University, Kasugai 487-8501, Japan; Graduate School of Bioagricultural Sciences, Nagoya University, Nagoya 464-8601, Japan

**Keywords:** Cytokinin receptor, Grafting, Root-derived cytokinin, *trans*-Zeatin

## Abstract

Compelling evidence demonstrates that root-derived cytokinins (CKs) contribute to shoot growth via long-distance transport; therefore, we hypothesized that an increase in root-derived CKs enhances shoot growth. To demonstrate this, we grafted Arabidopsis Col-0 (WT) scion onto rootstock originated from WT or a double-knockout line of CK receptors *AHK2* and *AHK3* (*ahk23*) because the knockout line over accumulates CKs in the body due to feedback homeostasis regulation. The grafted plants (scion/rootstock: WT/WT and WT/*ahk23*) were grown in vermiculite pots or solid media under high and low nitrogen regimes for vegetative growth and biochemical analysis. The root-specific deficiency of *AHK2* and *AHK3* increased root concentrations of *trans*-zeatin (tZ)-type and *N*^6^-(Δ^2^-isopentenyl) adenine (iP)-type CKs, induced CK biosynthesis genes, and repressed CK degradation genes in the root. Shoot growth, shoot concentrations of tZ-type CKs, and shoot expression of CK-inducible marker genes were consistently larger in the WT/*ahk23* plants than in the WT/WT plants. Moreover, the root-specific deficiency of *AHK2* and *AHK3* enhanced shoot growth in the WT scion more strongly than in the *ahk23* scion. Given that tZ-type CKs are predominantly produced from iP-type CKs in the root and xylem-mobile, it is concluded that the root-specific reduction of CK perception would enhance shoot growth by increasing the amount of root-derived tZ-type CKs and their perception by the shoot. This study will present a novel approach to improve plant growth and productivity.

## Introduction

Cytokinins (CKs) regulate various developmental processes and environmental responses, including cell division, shoot and root growth, cambial proliferation, and stress responses (Werner et al. 2001, Riefler et al. 2006, Sakakibara 2006, Matsumoto-Kitano et al. 2008). *N*^6^-(Δ^2^-isopentenyl) adenine (iP) and *trans*-zeatin (tZ) are biologically active forms of CKs in *Arabidopsis thaliana*. The initial rate-limiting step of their biosynthesis is the *N*^*6*^-prenylation of adenosine 5′-phosphates catalyzed by adenosine phosphate-isopentenyl transferases (IPTs), which produce iP-riboside 5′-phosphates (iPRPs) (Miyawaki et al. 2006, Osugi and Sakakibara 2015). Then, the iPRPs are converted into tZ-riboside 5′-phosphates (tZRPs) by two cytochrome P450 enzymes, CYP735A1, and CYP735A2 (Takei et al. 2004, Kiba et al. 2013). Finally, the iP-riboside 5′-monophosphate (iPRMP) and tZ-riboside (tZRMP) are dephosphoribosylated by the CK-activating enzyme LONELY GUY (LOG), which forms iP and tZ (Kurakawa et al. 2007, Kuroha et al. 2009, Tokunaga et al. 2012). They are perceived by three CK receptors, Arabidopsis HISTIDINE KINASE 2 (AHK2), AHK3, and AHK4/CRE1 (Inoue et al. 2001, Higuchi et al. 2004, Nishimura et al. 2004), activating the CK signaling cascade.

Compelling evidence indicates that CKs function as a long-distance signal transported through the vasculature (Sakakibara 2021). ATP-binding cassette transporter subfamily G14 (ABCG14) is expressed predominantly in the stele of roots and facilitates root-to-shoot transport of CKs (Ko et al. 2014, Zhang et al. 2014). Reciprocal grafting between wild-type and *abcg14* has demonstrated that CKs transported from roots contribute to shoot growth. The CYP735A1 and CYP735A2 are also predominantly expressed in roots (Takei et al. 2004), and thus, tZ-type CKs are biosynthesized mainly in the root. Shoot-specific impairment in the growth of *cyp735a1 chp735a2* mutants indicates that root-derived tZ-type CKs are crucial for shoot growth (Kiba et al. 2013). Grafting between various mutants on CK biosynthesis and transport mutants has demonstrated that root-derived tZ, a minor component of xylem CKs, controls leaf size but not the activity of the shoot apical meristem. By contrast, root-derived *trans*-zeatin riboside (tZR) contributes to both traits (Osugi et al. 2017). A recent study has reported that root-derived tZ-type CKs protect Arabidopsis shoots against photoperiod stress, abiotic stress caused by prolongation of the light period (Frank et al. 2020).

As mentioned above, root-derived CKs enhance growth, development, and stress tolerance in Arabidopsis shoots. This provoked us to produce plants whose shoots receive more CKs transported from roots, thereby showing better shoot growth. It is worthy of attention that deficiency of CK receptors significantly increases concentrations of tZ-type CKs in *A. thaliana*. By contrast, it impairs shoot growth because of a lack of CK perception in the shoot (Riefler et al. 2006). Thus, we supposed that root-specific reduction of CK perception could lead to CK accumulation in the root and facilitate their root-to-shoot transport, enhancing shoot growth. To demonstrate this, we grafted Arabidopsis Col-0 (WT) scion onto rootstock originated from WT or a double-knockout line of *AHK2* and *AHK3*. The grafted plants were cultivated to analyze their growth, CK concentrations, and expression of CK-related genes. Here we show that root-specific reduction of CK perception significantly enhances shoot growth and increases the levels of tZ-type CKs in the shoot. The possible underlying mechanisms are discussed.

## Results

### Confirmation of root-specific deficiency of AHK2 and AHK3

To achieve the root-specific reduction of CK perception, we aseptically grafted WT scion onto rootstock originated from WT or a double-knockout line of *AHK2* and *AHK3* using 6-day-old seedlings (Fig. 1A). We used *ahk2-5 ahk3-7* (*ahk23*) as the knockout line because this line contains the most tZ-type CKs among all double-knockout lines on *AHK2/3/4* and has vigorous roots for successful grafting (Riefler et al. 2006). At approximately 1 week after surgery, the grafted plants (scion/rootstock: WT/WT and WT/*ahk23*) having vigorous growth without any adventitious roots were selected for further experiments. The genomic polymerase chain reaction (PCR) and reverse transcription (RT)-PCR using the primers specific to *AHK2, AHK3*, T-DNA left border, and *ACTIN2* confirmed root-specific deficiency of *AHK2* and *AHK3* in the WT/*ahk23* plants (Fig. 1B, C).

**Fig. 1.**
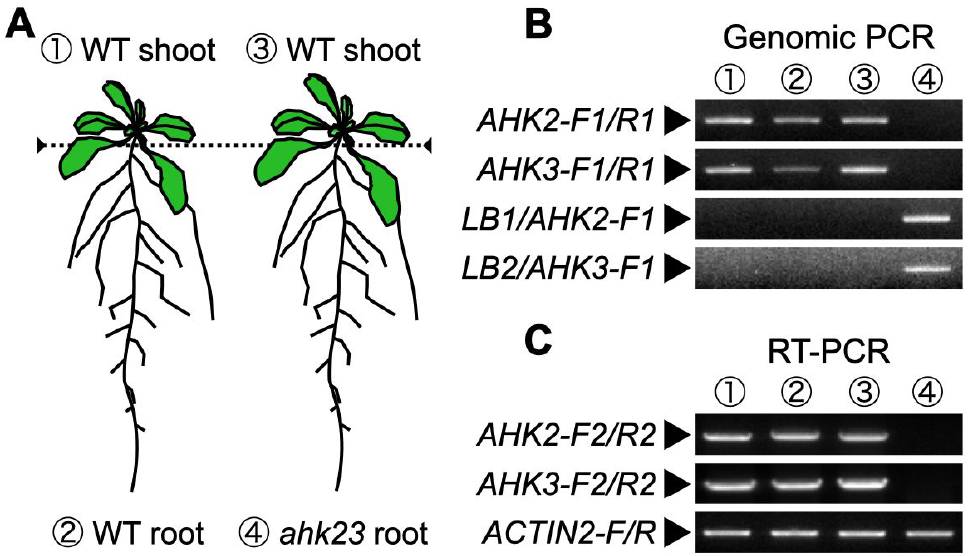
Root-specific deficiency of *AHK2* and *AHK3* by micrografting technique. (A) A schematic representation of two types of grafted plants. WT; wild-type Col-0, *ahk23*; *ahk2-5 ahk3-7* (Riefler et al. 2006). The dotted line represents the grafted seam in the hypocotyl. (B) A representative result of genomic PCR and agarose electrophoresis to confirm root-specific deficiency of *AHK2* and *AHK3. AHK2-F1, AHK2-R1, AHK3-F1, AHK3-R1, LB1*, and *LB2* are the specific primers used for PCR (Supplementary Table 1). (C) A representative result of RT-PCR and agarose electrophoresis to confirm root-specific deficiency of *AHK2* and *AHK3. AHK2-F2, AHK2-R2, AHK3-F2, AHK3-R2*, and *ACTIN2* are the specific primers used for PCR (Supplementary Table 1).

### Root-specific deficiency of AHK2 and AHK3 enhances shoot growth

The grafted plants were transferred to the vermiculite pots to analyze the effects of root-specific deficiency of CK perception on shoot growth. Then, the grafted plants were cultivated for 10 days under 10 mM nitrate (high N) or 0.5 mM nitrate (low N) because nitrogen availability is a strong determinant of CK contents in *A. thaliana* (Sakakibara et al. 2006). We observed their apparent shoot growth and determined the fresh weight, dry weight, rosette diameter, leaf area, and leaf number in their shoots (Fig. 2A–F). The WT/*ahk23* plants had larger shoot growth than did the WT/WT plants under both nitrate conditions (Fig. 2A). The fresh shoot weight and dry weight were greater in the WT/*ahk23* plants than in the WT/WT plants by 21% and 29% under high N and by 18% and 35% under low N on average, respectively (Fig. 2B, C). There were significant increases in the rosette diameter and leaf area in the WT/*ahk23* plants relative to the WT/WT plants (Fig. 2D, E). Alternatively, the leaf number was comparable between the two types of grafted plants (Fig. 2F). *In vitro* culture of the grafted plants on the solid media also confirmed that fresh shoot weight was significantly larger in the WT/*ahk23* plants than in the WT/WT plants by 18% under high N and 20% under low N averagely (Supplementary Fig. S1A).

**Fig. 2.**
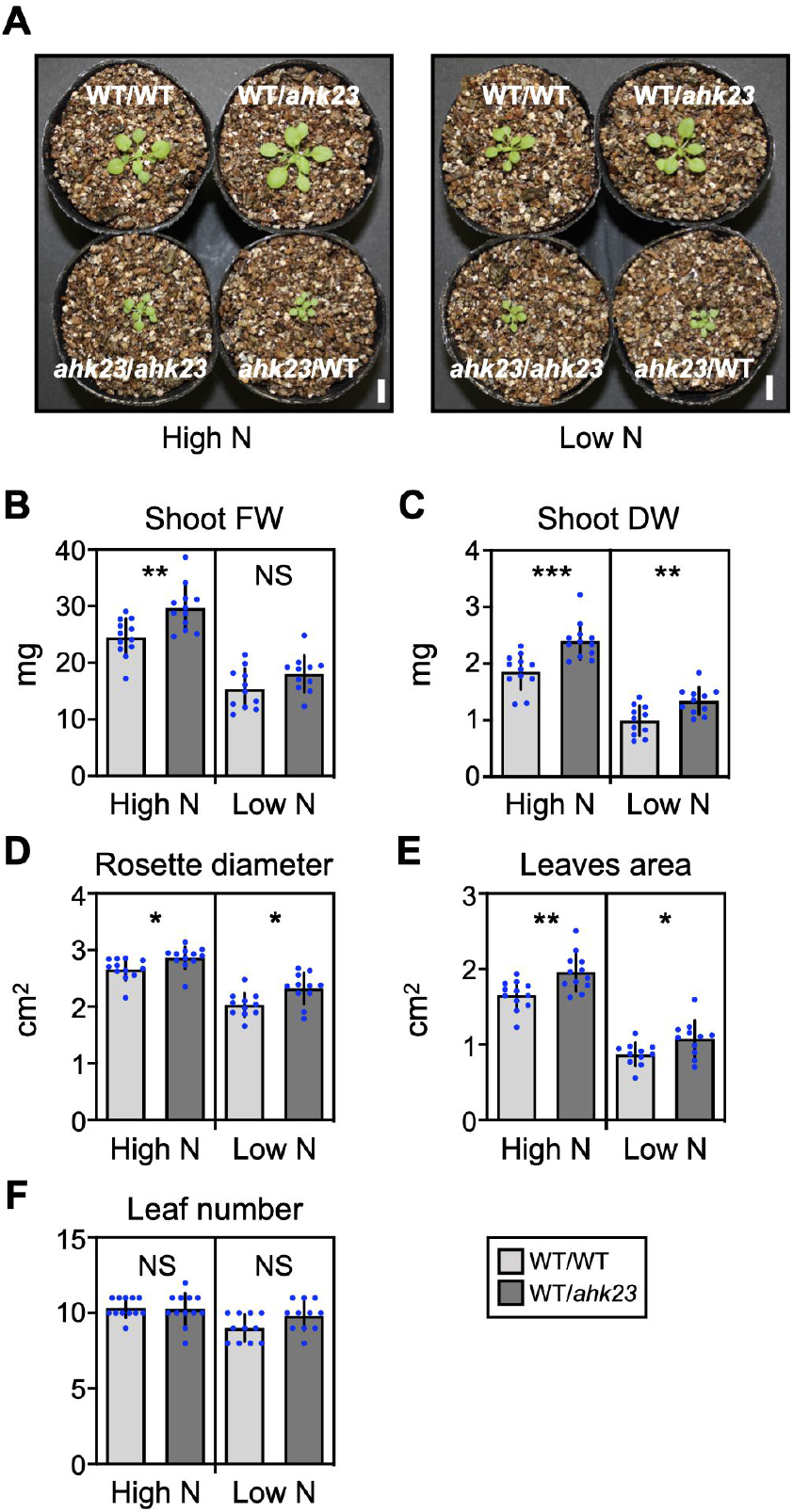
Effects of root-specific deficiency of *AHK2* and *AHK3* on shoot growth. The grafted plants were transferred to the vermiculite pots and grown for 10 days with a supply of MGRL-based salts containing 10 mM (high N) or 0.5 mM (low N) nitrate. The shoot appearance (A), shoot fresh weight (B), shoot dry weight (C), rosette diameter (D), leaf area (E), and leaf number (F) of the grafted plants were analyzed. Data are presented as the mean ± SD (n = 11–12). WT/WT, WT/*ahk23, ahk23*/WT, and *ahk23*/*ahk23* denote the plants corresponding to the WT scion, WT scion, *ahk23* scion, and *ahk23* scion grafted on the WT rootstock, *ahk23* rootstock, WT rootstock, and *ahk23* rootstock, respectively. \**P <* 0.05; \*\**P <* 0.01; \*\*\**P <* 0.001 (Welch’s *t*-test). NS denotes not significant. FW, fresh weight; DW, dry weight. The scale bar denotes 1 cm.

### Root-specific deficiency of AHK2 and AHK3 does not affect the root growth

It has been widely reported that the length and number of lateral roots are increased by a deficiency of CK biosynthesis, perception, and signaling (Miyawaki et al. 2006, Riefler et al. 2006). However, in this study, no significant difference was observed in the total root length and lateral root number between the WT/WT and WT/*ahk23* plants vertically grown on solid media containing 10 or 0.5 mM nitrate (Fig. 3A, B). Also, the fresh root weight was comparable between the horizontally grown grafted plants (Supplementary Fig. S1B).

**Fig. 3.**
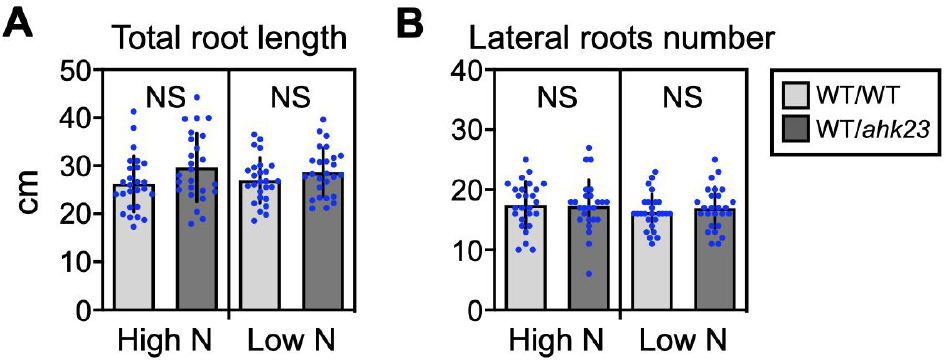
Effects of root-specific deficiency of *AHK2* and *AHK3* on root growth. The grafted plants were transferred to nitrogen-modified 1/2MS media containing 10 mM (high N) or 0.5 mM (low N) nitrate and grown vertically for 3 days. The total root length (A) and lateral root number (B) of the grafted plants were analyzed. Quantitative data are presented as the mean ± SD (n = 25–26). NS denotes not significant (Welch’s *t*-test).

### Root-specific deficiency of AHK2 and AHK3 increases root tZ-type CK levels, induces CK biosynthesis genes, and represses CK degradation genes

Next, to elucidate the effects of root-specific deficiency of CK perception on the root CK levels, we determined tZ-type and iP-type CKs in the root of grafted plants. Not only concentrations of tZ and tZR but also those of the other tZ-type CKs and iP-type CKs were consistently higher in the WT/*ahk23* roots than in the WT/WT roots under both nitrate regimes (Fig. 4A–E and Supplementary Fig. S2A–D). The tZ and tZR concentrations in the WT/*ahk23* roots were averagely 2.8 and 1.9 times under high N and 2.7 and 2.3 times under low N as much as those in the WT/WT roots, respectively (Fig. 4A, B). The concentrations of respective CK species were generally larger under the 10 mM nitrate conditions than under the 0.5 mM nitrate conditions (Fig. 4A–E and Supplementary Fig. S2A–D).

**Fig. 4.**
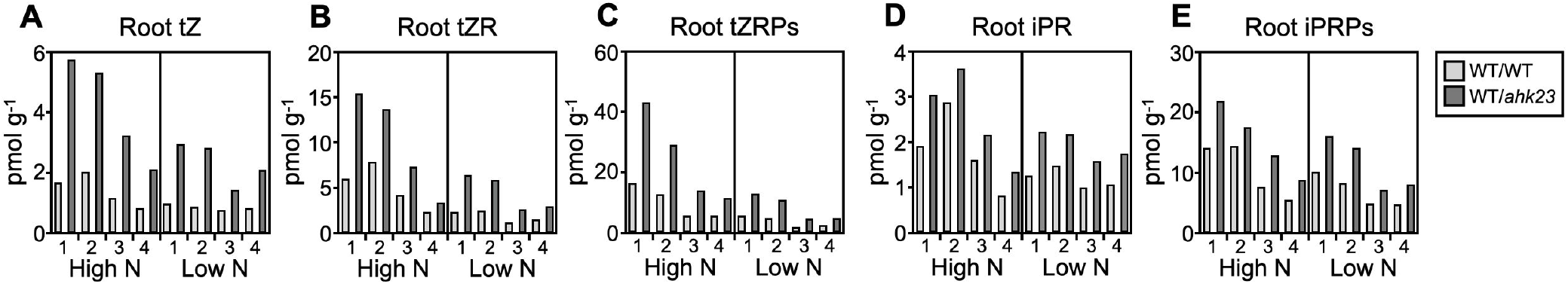
Effects of root-specific deficiency of *AHK2* and *AHK3* on cytokinin concentrations in the root. The grafted plants were transferred to nitrogen-modified 1/2MS media containing 10 mM (high N) or 0.5 mM (low N) nitrate and grown in a horizontal position for 7 days. The samples harvested from four independent grafting experiments were subjected to cytokinin analysis. ‘1’, ‘2’, ‘3’, and ‘4’ below the graph mean the 1st, 2nd, 3rd, and 4th grafting experiment, respectively. Roots from five plants were pooled as one biological replicate. The cytokinin analysis for the 1st and 2nd samples and that for the 3rd and 4th samples were separately performed. The root concentrations of tZ (A), tZR (B), tZRPs (C), iPR (D), and iPRPs (E) in the grafted plants are shown. tZ, *trans*-zeatin; tZR, tZ-riboside; tZRPs, tZR 5′-phosphates; iPR, *N*^*6*^-(Δ^2^-isopentenyl)adenine riboside; iPRPs, iPR 5′-phosphates.

To analyze how root-specific deficiency of CK perception increases the root CK levels, we performed quantitative RT-PCR analysis on the genes of the *CYTOKININ OXIDASE/DEHYDROGENASEs* (*CKXs*) involved in CK degradation and the *IPTs* and *CYP735A1/A2* contributing to *de novo* biosynthesis of iP-type CKs and their conversion to tZ-type CKs. While the expression of *CKXs* except for *CKX3* was downregulated by root-specific deficiency of *AHK2* and *AHK3*, that of *IPT1, IPT3, IPT5*, and *IPT7* was upregulated (Fig. 5). Notably, the transcript levels of *CKX2* and *CKX4* were lower in the WT/*ahk23* roots than in the WT/WT roots by 93% and 91% under high N and 95% and 84% under low N on average, respectively (Fig. 5). The opposite changes in *IPTs* and *CKXs* expression would collectively enhance CK accumulation in roots deficient in CK perception. The root expression of *CYP735A2* was comparable between the grafted plants under high N conditions, whereas that was larger in the WT/*ahk23* plants than in the WT/WT plants under low N conditions (Fig. 5). Irrespective of N conditions, the expression of *CYP735A1*, a minor isogene, was higher in the WT/*ahk23* roots than in the WT/WT roots (Fig. 5). These suggest that root-specific deficiency of CK perception could upregulate the root genes contributing to tZ-type CK biosynthesis, especially under low N conditions. The expression of *ABCG14* involved in the root-to-shoot transport of tZ-type CKs was little changed depending on the grafting (Fig. 5).

**Fig. 5.**
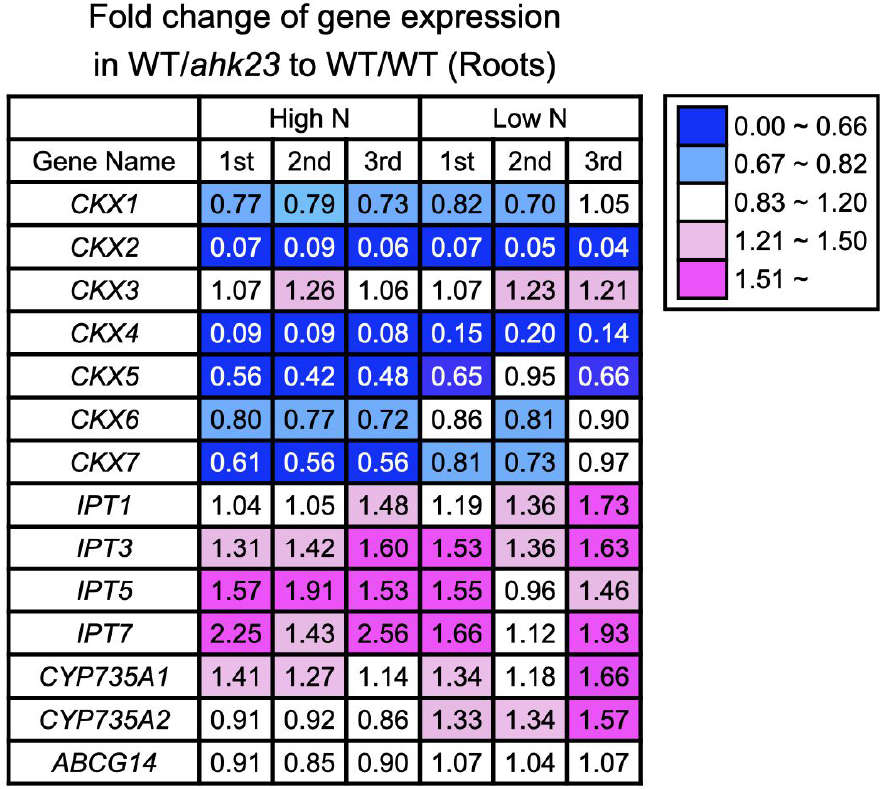
Effects of root-specific deficiency of *AHK2* and *AHK3* on the expression of genes contributing to cytokinin degradation, biosynthesis, and transport in the root. The grafted plants were transferred to nitrogen-modified 1/2MS media containing 10 mM (high N) or 0.5 mM (low N) nitrate and grown in a horizontal position for 7 days. The samples harvested from three independent grafting experiments were subjected to total RNA purification and subsequent RT-qPCR. ‘1st’, ‘2nd’, and ‘3rd’ mean the 1st, 2nd, and 3rd grafting experiment, respectively. Roots from five plants were pooled as one biological replicate. The relative transcript levels in the WT/*ahk23* roots are shown regarding those in the WT/WT roots as one.

### Root-specific deficiency of AHK2 and AHK3 increases tZ-type CK levels and expression of CK-inducible genes in the shoot

To analyze the effects of root-specific deficiency of CK perception on the shoot CK levels, we determined tZ-type and iP-type CKs in the shoots of grafted plants. Shoot concentrations of tZ-type CKs were consistently higher in the WT/*ahk23* plants (Fig. 6A–C and Supplementary Fig. S3A, B). The tZ and tZR concentrations in the WT/*ahk23* shoots were averagely 1.4 and 1.7 times under high N and 1.4 and 1.5 times under low N as much as those in the WT/WT shoots, respectively (Fig. 6A, B). By contrast, shoot concentrations of iP-type CKs were not always higher in the WT/*ahk23* plants than in the WT/WT plants (Fig. 6D, E and Supplementary Fig. S3C, D). These imply that the root-to-shoot transport of xylem-mobile tZ and tZR could be larger in the WT/*ahk23* plants than in the WT/WT plants. The shoot concentrations of tZ, tZR, tZRPs, iPR, and iPRPs were generally larger under high N than under low N in the same way as the roots (Fig. 6A–E), whereas those of cytokinin *N*- and *O*-glucosides were similar under both N conditions (Supplementary Fig. S3A, B, D).

**Fig. 6.**
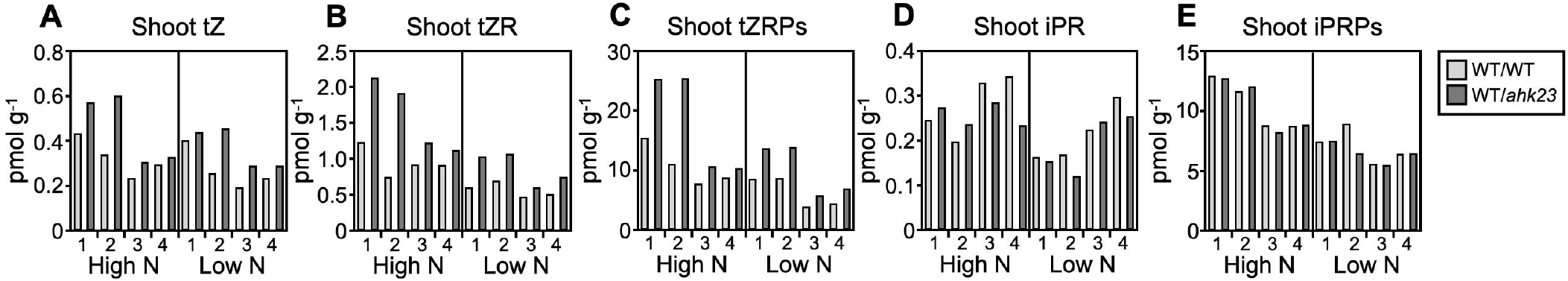
Effects of root-specific deficiency of *AHK2* and *AHK3* on cytokinin concentrations in the shoot. The grafted plants were transferred to nitrogen-modified 1/2MS media containing 10 mM (high N) or 0.5 mM (low N) nitrate and grown in a horizontal position for 7 days. The samples harvested from four independent grafting experiments were subjected to cytokinin analysis. ‘1’, ‘2’, ‘3’, and ‘4’ below the graph mean the 1st, 2nd, 3rd, and 4th grafting experiment, respectively. Shoots from five plants were pooled as one biological replicate. The cytokinin analysis for the 1st and 2nd samples and that for the 3rd and 4th samples were separately performed. The shoot concentrations of tZ (A), tZR (B), tZRPs (C), iPR (D), and iPRPs (E) in the grafted plants are shown. tZ, *trans*-zeatin; tZR, tZ-riboside; tZRPs, tZR 5′-phosphates; iPR, *N*^*6*^-(Δ^2^-isopentenyl) adenine riboside; iPRPs, iPR 5′-phosphates.

To verify whether the increased tZ-type CKs stimulate CK signaling in the WT/*ahk23* shoots, we checked the gene expression of CK-inducible type-A *ARABIDOPSIS RESPONSE REGULATORS* (*ARRs*) (Osugi et al. 2017). The expression of the *ARRs* was upregulated in the WT/*ahk23* shoots compared with the WT/WT shoots, and upregulation was pronounced under low N conditions compared with high N conditions (Fig. 7). This suggests that root-specific deficiency of CK perception would enhance CK signaling in the shoot.

**Fig. 7.**
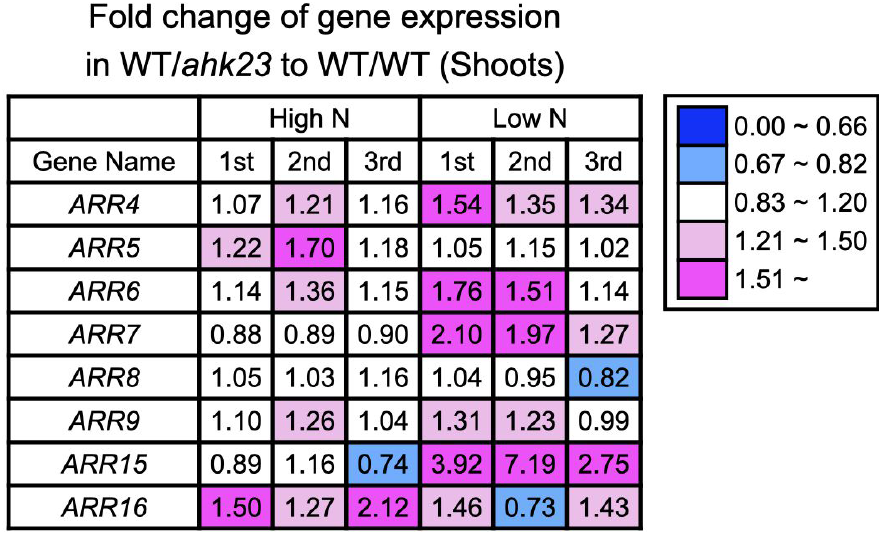
Effects of root-specific deficiency of *AHK2* and *AHK3* on the expression of cytokinin-inducible type-A *ARR* genes in the shoot. The grafted plants were transferred to nitrogen-modified 1/2MS media containing 10 mM (high N) or 0.5 mM (low N) nitrate and grown in a horizontal position for 7 days. The samples harvested from three independent grafting experiments were subjected to total RNA purification and subsequent RT-qPCR. 1st’, ‘2nd’, and ‘3rd’ mean the 1st, 2nd, and 3rd grafting experiment, respectively. Shoots from five plants were pooled as one biological replicate. The relative transcript levels in the WT/*ahk23* shoots are shown regarding those in the WT/WT shoots as one.

## Discussion

CK signaling enhances the growth and development of shoots, whereas negatively regulates those of roots (Miyawaki et al. 2006, Riefler et al. 2006, Osugi et al. 2017). Consequently, the deficiency of genes for CK biosynthesis, perception, and signaling leads to relatively smaller shoots and larger roots. Interestingly, root-specific reduction of CK contents by constitutively expressing *CKX* under the control of root-specific promoter enhances root growth and mineral absorption efficiency apparently without impairing vegetative shoot growth in *A. thaliana* and tobacco (Werner et al. 2010). Similar approaches have succeeded in improving root growth and crop yields in barley and chickpea (Ramireddy et al. 2018, Khandal et al. 2020). On the other hand, root-specific expression of *CKX* significantly decreases the shoot concentrations of tZ-type and iP-type CKs, albeit with no negative effects on shoot growth (Werner et al. 2010). Since the tZ-type CKs transported from roots promote shoot growth (Kiba et al. 2013, Osugi et al. 2017), this study aimed to increase the amounts of root-derived CKs by root-specific reduction of CK perception. Expectedly, CKs accumulated in the roots lacking AHK2 and AHK3 (Fig. 4 and Supplementary Fig. S2), probably because of the induction of *IPTs* and repression of *CKXs* (Fig. 5). Shoot growth, shoot concentrations of tZ-type CKs, and shoot expression of CK-inducible marker genes were consistently increased in the WT/*ahk23* plants compared with that in the WT/WT (Fig. 2, 6A–C, 7, and Supplementary Fig. S1A, S3A, B). Previous studies have demonstrated that tZ-type CKs are predominantly produced from iP-type CKs in the root (Kiba et al. 2013) and tZ and tZR are xylem-mobile from root-to-shoot (Osugi et al. 2017). Also, we observed that the shoot expression of *CYP735A2* was approximately 1% as much as its root expression and was not significantly induced by the root-specific deficiency of *AHK2* and *AHK3* (Supplementary Fig. S5). The *CYP735A1* expression was not detected in the shoot (data not shown). These indicate that the increased tZ-type CKs in the WT/*ahk23* shoots could originate from their roots. Kiba et al. (2013) have reported that *cyp735a1 cyp735a2* mutants lacking the tZ-type CKs grew rosette leaves with reduced leaf blade size and petiole length compared with WT plants, whereas their leaf number was not altered. In this study, root-specific reduction of CK perception increased leaf area and rosette diameter but not leaf number (Fig. 2), which indicates the increased action of tZ-type CKs. Moreover, we observed that the root-specific deficiency of *AHK2* and *AHK3* enhanced shoot growth in the WT scion more effectively than in the *ahk23* scion except for the rosette diameter under high N (Fig. 2A and Supplementary Fig. S4), suggesting that a significant part of shoot growth enhancement would occur through increased shoot perception of CKs. Altogether, we conclude that the root-specific reduction of CK perception would enhance shoot growth by increasing the amounts of root-derived tZ-type CKs and their perception by the shoot.

The root-specific deficiency of *AHK2* and *AHK3* induced the shoot expression of CKs-inducible *ARRs* more intensively under low N than under high N except for *ARR5* and *ARR16* (Fig. 7). The basal levels of tZ, tZR, and tZRPs in shoots were generally lower under low N than under high N (Fig. 6A–C). Hence, the increase in root-derived CKs could drive the CK signaling more effectively in the shoots lacking CKs than those filled with CKs. On the other hand, the root-specific *AHK2* and *AHK3* deficiency enhanced shoot growth almost equally between the two N regimes (Fig. 2 and Supplementary Fig. S1). Thus, in the shoot, the increase in CK signaling is not always correlated with growth enhancement, implying that the unknown signaling molecule(s) other than CKs might be also involved in the growth enhancement.

Kiba et al. (2011) reported that under low N conditions, exogenous application of tZ significantly represses the expression of high-affinity nitrate transporters *NRT2*.*1* and *NRT2*.*4*, depending on the functional AHK3 and AHK4/CRE1. This implies that root-specific reduction of CK perception could elevate *NRT2*.*1* and *NRT2*.*4* expression, thereby facilitating nitrate acquisition, especially under low N conditions. However, the transcript levels of *NRT2*.*1* and *NRT2*.*4* were not significantly different between the roots of WT/WT and WT/*ahk23* regardless of the N regime (Supplementary Fig. S6A, B). Also, the nitrate concentrations in the shoot and the root did not significantly differ between the grafted plants (Supplementary Fig. S6C, D). It awaits further challenges by the root-specific deficiency of *AHK3* and *AHK4* or *AHK2* and *AHK3* and *AHK4* to improve N uptake ability.

Although root-specific reduction of CK contents strongly enhances root growth and development in previous studies (Werner et al. 2010, Ramireddy et al. 2018, Khandal et al. 2020), our root-specific reduction of CK perception did not significantly alter them (Fig. 3 and Supplementary Fig. S1B). In this study, the plants before and after grafting were cultivated under relatively low-light conditions (photosynthetic photon flux density [PPFD]: approximately 40–50 μmol m^-2^ s^-1^) to alleviate photooxidative stress under low N conditions (Cetner et al. 2017). Therefore, the shortage of carbon sources and additional carbon consumption by shoot growth enhancement might limit the biomass allocation to the WT/*ahk23* root. In the future study, we will test whether higher light intensities or carbon dioxide concentrations improve root growth.

In the present study, we have succeeded in increasing shoot growth by root-specific reduction of CK perception. Nowadays, the genome editing technologies are applicable for disrupting a gene of interest in various plant species and therefore, the rootstocks lacking CK receptors would be available in useful crops. This study presents a novel approach to improve crop growth and productivity.

## Materials and Methods

### Plant materials and growth conditions

Seeds of Col-0 and *ahk2-5 ahk3-7* (Riefler et al. 2006) were surface-sterilized and sown in the solid medium containing half-strength Murashige and Skoog (1/2MS) salts, supplemented with 0.05% (w/v) MES-KOH (pH 5.7), 1% (w/v) sucrose, and 0.5% (w/v) gellan gum (Fujifilm Wako, Osaka, Japan). The seeds were placed in the dark at 4°C for 3–5 days to break dormancy. Seedlings were grown vertically under a PPFD of 25 μmol m^-2^ s^-1^ (16 h light/8 h dark cycle) at 22°C. The micrografting was aseptically performed using the 6-day-old seedlings. The seedlings were cut perpendicularly in the hypocotyl with an injection needle tip (Code NN-2613S; TERUMO, Tokyo, Japan). The obtained scion was connected with the partner rootstock in a silicon microtube (Code 1-8194-04; AS ONE, Osaka, Japan). The grafted plants were incubated for 5 days under a PPDF of approximately 40–50 μmol m^-2^ s^-1^ (constant light) at 27°C and further cultivated for a few days under a PPFD of approximately 25 μmol m^-2^ s^-1^ (16 h light/8 h dark cycle) at 22°C. The successfully grafted plants were selected, transferred to vermiculite pots or solid media, and grown for the following experiments. For vermiculite cultivation, the plants were grown under a PPFD of approximately 40–50 μmol m^-2^ s^-1^ (16 h light/8 h dark cycle) at 22°C. The 30 mL of nitrogen-modified MGRL-based salts (Fujiwara et al. 1992); including 10 mM potassium nitrate (high N) or 0.5 mM potassium nitrate and 9.5 mM potassium chloride (low N), was supplied twice before harvest. For the *in vitro* culture on the solid media, the plants were grown under a PPFD of approximately 40– 50 μmol m^-2^ s^-1^ (constant light) at 22°C in the nitrogen-modified 1/2MS salts including 10 mM potassium nitrate (high N) or 0.5 mM potassium nitrate and 9.5 mM potassium chloride (low N) supplemented with 0.1% (w/v) MES-KOH (pH 5.7), 1% (w/v) sucrose, and 0.25% (horizontally) or 0.5% (vertically) (w/v) gellan gum (Wako, Osaka, Japan). To allow nondestructive sampling of the roots in horizontally grown plants, the surfaces of solid media were covered with cellophane sheets (Hachiya et al. 2021). The other details are described in the figure legends.

### Extraction of total RNA

Shoots and roots were harvested, immediately frozen with liquid N_2_, and stored at −80°C until use. Shoots and roots from five plants were pooled as one biological replicate. Frozen samples were ground with a Multi-Beads Shocker (Yasui Kikai Corp., Osaka Prefecture, Osaka, Japan) using zirconia beads (diameter, 5 mm). Total RNA was extracted using the RNeasy Plant Mini Kit (Qiagen Japan, Tokyo, Japan) according to the manufacturer’s instructions.

### Quantitative RT-PCR

Reverse transcription was performed using a ReverTra Ace qPCR RT Master Mix with gDNA Remover (Toyobo Co. Ltd., Tokyo, Japan) according to the manufacturer’s instructions. The synthesized cDNA was diluted fivefold with distilled water and used in quantitative PCR (qPCR). Transcript levels were measured using a TaKaRa Thermal Cycler Dice TP800 (TaKaRa Bio Inc., Shiga, Japan). The obtained cDNA (2 µL) was amplified in the presence of 10 µL KOD SYBR qPCR Mix (TOYOBO Co. Ltd.), 0.4 µL specific primers (0.2 µM final concentration), and 7.2 µL sterile distilled water. Relative transcript levels were calculated using the comparative cycle threshold (Ct) method using *ACTIN3* as the internal standard (Hachiya et al. 2020). The primer sequences used in the experiments are shown in Supplementary Table S1.

### Determination of CK species

Shoots and roots were snap-frozen with liquid N_2_ and stored at −80°C until use. Shoots and roots from five plants were pooled as one biological replicate. Extraction and determination of CKs were performed as described previously (Kojima et al. 2009); using ultra-performance liquid chromatography (UPLC)– tandem quadrupole mass spectrometry (AQUITY UPLC System/XEVO-TQS; Waters) with an octadecylsilyl column (ACQUITY UPLC HSS T3, 1.8 µm 2.1 mm × 100 mm).

### Determination of nitrate

Nitrate was determined according to the method reported by Hachiya and Okamoto (2017). Shoots and roots were snap-frozen with liquid N_2_ and stored at −80°C until use. Shoots and roots from independent plates were pooled separately as single biological replicates for samples. The shoots and roots were boiled with 10 volumes of H_2_O at 100°C for 20 min. The obtained 10 μL supernatant or the 10 μL standard solution of potassium nitrate was mixed with a 40 μL reaction reagent containing 50 mg salicylic acid per 1 mL sulfuric acid, followed by incubation at room temperature for 20 min. For the mock preparation, 40 μL sulfuric acid alone instead of the reaction reagent was added to the 10 μL supernatant. After adding 1 mL of 8% (w/v) NaOH solution to the mixture, the absorbance at 410 nm was scanned using a plate reader (CORONA SH-9000Lab, Hitachi High-Tech Corp., Tokyo, Japan). The nitrate concentration was calculated from the standard curve.

### Statistical

Statistical analyses were done using the R software v.2.15.3.

## Acknowledgments

The authors would like to thank Enago (www.enago.jp) for the English language review. Seeds of *ahk2-5 ahk3-7* were kindly provided by Prof. Thomas Schmülling. We thank Mr. Takahiro Oya (Shimane Univ.), Ms. Natsumi Ooi (Shimane Univ.), and Mr. Kosuke Yoshizawa (Shimane Univ.) for their technical assistance.

