## Supplemental Data for "Root-specific reduction of cytokinin perception enhances shoot growth with an alteration of *trans*-zeatin distribution in *Arabidopsis thaliana*"

**Supplementary Figure S1**

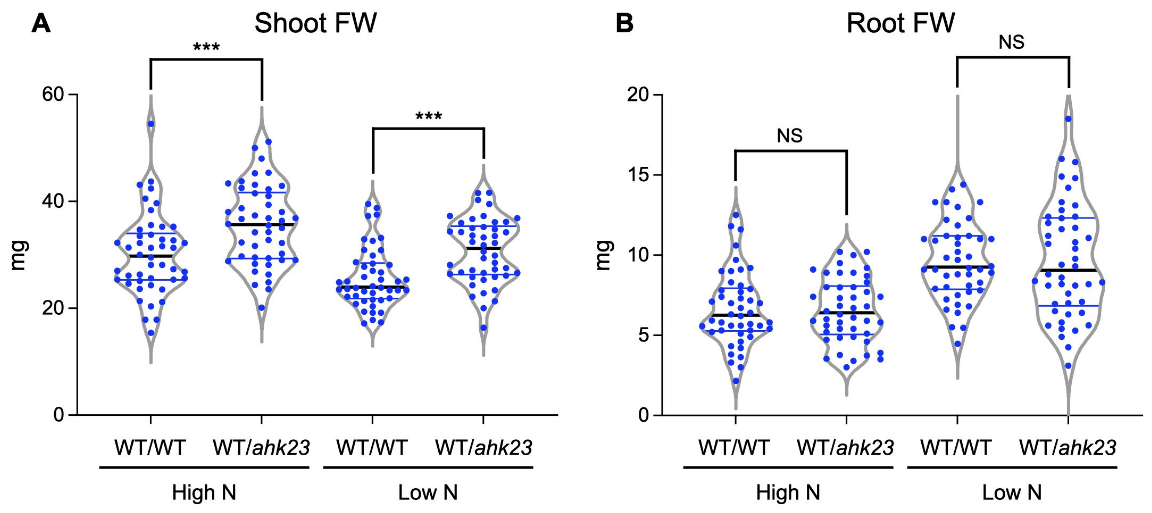


**Fig. S1 Violin plot of the fresh weights in shoots and roots of the grafted plants**

The grafted plants were transferred to nitrogen-modified 1/2MS media containing 10 mM (high N) or 0.5 mM (low N) nitrate and horizontally grown for 7 days. All the data pooled from six independent grafting experiments are shown (n = 46). **P* < 0.05; ***P* < 0.01; ****P* < 0.001 (Welch's *t*-test). NS denotes not significant. The central thick solid line and two thin solid lines represent the median and quartiles, respectively.

### Supplementary Figure S2

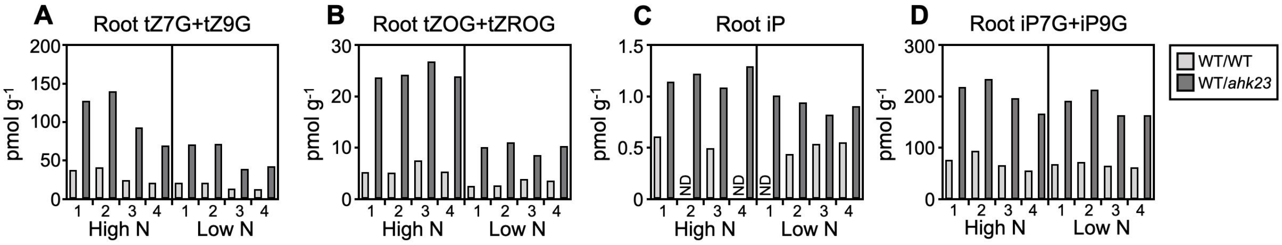


**Fig. S2 Effects of root-specific deficiency of *AHK2* and *AHK3* on cytokinin concentrations in the root**

The grafted plants were transferred to nitrogen-modified 1/2MS media containing 10 mM (high N) or 0.5 mM (low N) nitrate and grown in a horizontal position for 7 days. The samples harvested from four independent grafting experiments were subjected to cytokinin analysis. '1', '2', '3', and '4' below the graph mean the 1st, 2nd, 3rd, and 4th grafting experiment, respectively. Roots from five plants were pooled as one biological replicate. The cytokinin analysis for the 1st and 2nd samples and that for the 3rd and 4th samples were separately performed. The root concentrations of t tZ7G + tZ9G (A), tZOG + tZROG (B), iP (C), and iP7G + 9G (D) in the grafted plants are shown. tZ7G, tZ-7-*N*-glucoside; tZ9G, tZ-9-*N*-glucoside; tZOG, tZ-*O*-glucoside; tZROG, tZR-*O*-glucoside; iP7G, iP-7-*N*-glucoside; iP9G, iP-9-*N*-glucoside. ND means 'not detected'.

### Supplementary Figure S3

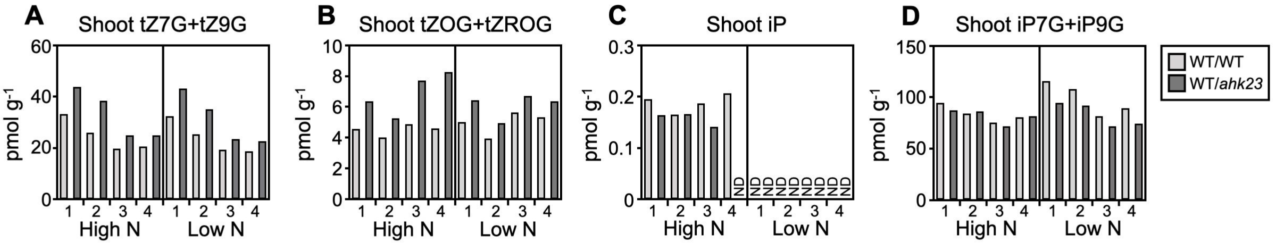


**Fig. S3 Effects of root-specific deficiency of *AHK2* and *AHK3* on cytokinin concentrations in the shoot**

The grafted plants were transferred to nitrogen-modified 1/2MS media containing 10 mM (high N) or 0.5 mM (low N) nitrate and grown in a horizontal position for 7 days. The samples harvested from four independent grafting experiments were subjected to cytokinin analysis. '1', '2', '3', and '4' below the graph mean the 1st, 2nd, 3rd, and 4th experiment, respectively. Shoots from five plants were pooled as one biological replicate. The cytokinin analysis for the 1st and 2nd samples and that for the 3rd and 4th samples were separately performed. The shoot concentrations of tZ7G + tZ9G (A), tZOG + tZROG (B), iP (C), and iP7G + iP9G (D) in the grafted plants are shown. tZ7G, tZ-7-*N*-glucoside; tZ9G, tZ-9-*N*-glucoside; tZOG, tZ-*O*-glucoside; tZROG, tZR-*O*-glucoside; iP7G, iP-7-*N*-glucoside; iP9G, iP-9-*N*-glucoside. ND means 'not detected'.

### Supplementary Figure S4

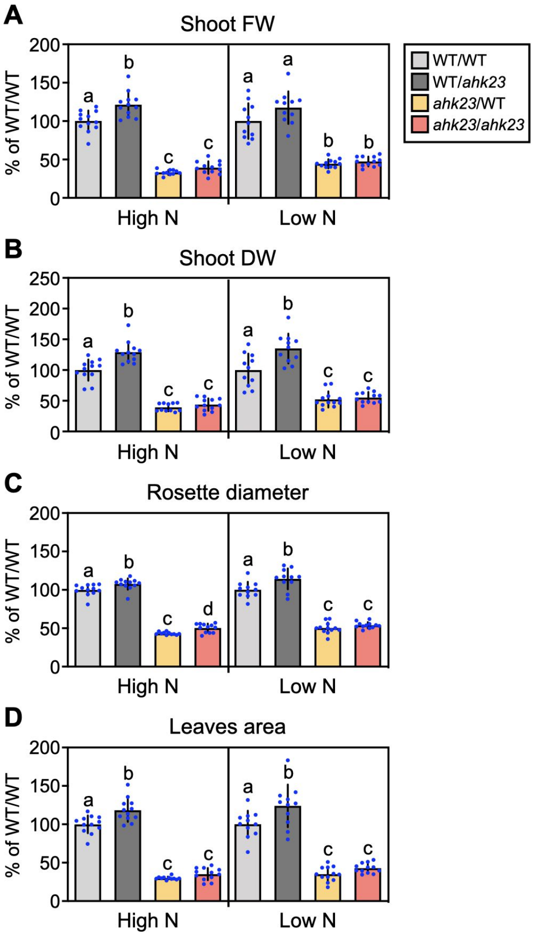


**Fig. S4 Effects of root-specific deficiency of *AHK2* and *AHK3* on shoot growth**

The grafted plants were transferred to the vermiculite pots and grown for 10 days with a supply of MGRL-based salts containing 10 mM (high N) or 0.5 mM (low N) nitrate. The fresh shoot weight (A), shoot dry weight (B), rosette diameter (C), and leaf area (D) of the grafted plants were analyzed. Data are presented as the mean ± SD (n = 11–12). WT/WT, WT/*ahk23*, *ahk23*/WT, and *ahk23*/*ahk23* denote the plants corresponding to the WT scion, WT scion, *ahk23* scion, and *ahk23* scion grafted on the WT rootstock, *ahk23* rootstock, WT rootstock, and *ahk23* rootstock, respectively. Different lowercase letters indicate significant differences evaluated by the Tukey–Kramer multiple comparison tests conducted at a significance level of $P < 0.05$. FW, fresh weight; DW, dry weight.

**Supplementary Figure S5**

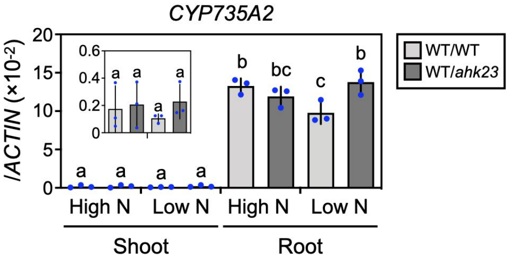


**Fig. S5 Effects of root-specific deficiency of *AHK2* and *AHK3* on the expression of *CYP735A2* in the shoot and root**

The grafted plants were transferred to nitrogen-modified 1/2MS media containing 10 mM (high N) or 0.5 mM (low N) nitrate and grown in a horizontal position for 7 days. The samples harvested from three independent grafting experiments were subjected to total RNA purification and subsequent RT-qPCR. Data are presented as the mean ± SD (n = 3). Different lowercase letters indicate significant differences evaluated by the Tukey–Kramer multiple comparison tests conducted at a significance level of $P < 0.05$. The inset is the magnification corresponding to the shoot expression of *CYP735A2*.

### Supplementary Figure S6

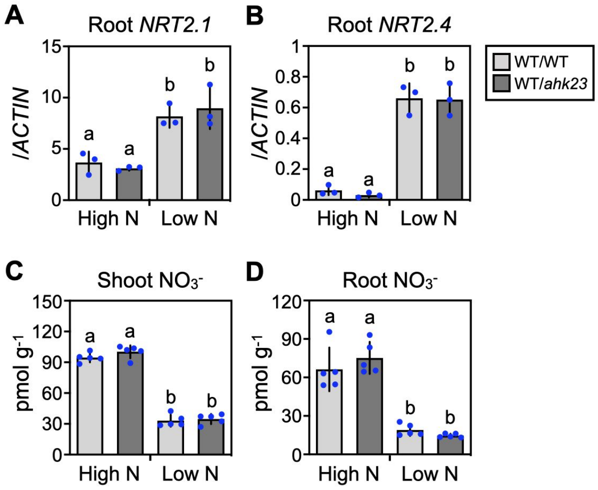


**Fig. S6 Effects of root-specific deficiency of *AHK2* and *AHK3* on root expression of *NRT2.1* and *NRT2.4* and nitrate concentrations in the shoot and root**

The grafted plants were transferred to nitrogen-modified 1/2MS media containing 10 mM (high N) or 0.5 mM (low N) nitrate and grown in a horizontal position for 7 days. The harvested samples were subjected to total RNA purification and subsequent RT-qPCR and nitrate determination. The transcript levels of *NRT2.1* (A) and *NRT2.4* (B) in the root are presented as the mean ± SD (n = 3). The nitrate concentrations in the shoot (C) and root (D) are shown as the mean ± SD (n = 5). Different lowercase letters indicate significant differences evaluated by the Tukey–Kramer multiple comparison tests conducted at a significance level of $P < 0.05$.

**Supplementary Table S1. Primers used in this study**

| Primer name | Purpose | Primer sequences |
|---|---|---|
| *AHK2-F1* | Genotyping | GCAAGAGGCTTTAGCTCCAA |
| *AHK2-R1* | Genotyping | TTGCCCGTAAGATGTTTTCA |
| *AHK3-F1* | Genotyping | AATTTCAGCCTTTGGGTTGTC |
| *AHK3-R1* | Genotyping | CTGAATGGAGCACCCTCATAG |
| *LB1* | Genotyping | GCCTTTTCAGAAATGGATAAATAGCCTTGCTTCC |
| *LB2* | Genotyping | ATATTGACCATCATACTCATTGC |
| *ABCG14-Q-F* | RT-qPCR | ATCTGTTACTACACTCGTTTTTCTC |
| *ABCG14-Q-R* | RT-qPCR | TGTAGCAGTAGTAGCTGTAGCTTAG |
| *ACTIN2-F* | RT-qPCR | TGTCCTCCTCACTTTCATCAGC |
| *ACTIN2-R* | RT-qPCR | CATCAATTCGATCACTCAGAGC |
| *ACTIN3-Q-F* | RT-qPCR | GGCTAACCGTGAGAAGATGA |
| *ACTIN3-Q-R* | RT-qPCR | CGACCTGCAAGATCAAGACG |
| *AHK2-RT-F2* | RT-qPCR | AGTCGTCGAGCAGTGACAAG |
| *AHK2-RT-R2* | RT-qPCR | TTGCTACCGCTGTGTAGAGC |
| *AHK3-RT-F2* | RT-qPCR | TCTGGATGCTATGCTGCTGG |
| *AHK3-RT-R2* | RT-qPCR | CCTGTACAGCTGCTCTGCTT |
| *ARR4-Q-F* | RT-qPCR | GTTGACTGTTTCGACTGAATC |
| *ARR4-Q-R* | RT-qPCR | GTCATCATCTTCATCGTCTATC |
| *ARR5-Q-F* | RT-qPCR | GCTGTTGATGATAGTATGGTTG |
| *ARR5-Q-R* | RT-qPCR | AGATATTGTAAAGCTCTTGTCG |
| *ARR6-Q-F* | RT-qPCR | GAAGTTATGCTACCGAGGAAG |
| *ARR6-Q-R* | RT-qPCR | TACGATCAACGTGACTGTCGT |
| *ARR7-Q-F* | RT-qPCR | AAGCCATTCTAACAAGAGAAAG |
| *ARR7-Q-R* | RT-qPCR | GAACATGAAGAGTCCTTGATAG |
| *ARR8-Q-F* | RT-qPCR | GTGTCTAAACCGGAGATAGAAG |
| *ARR8-Q-R* | RT-qPCR | ACTACTCAACATTGGTTCAAGT |
| *ARR9-Q-F* | RT-qPCR | TGCAACAAGATCTGCTATTAGT |
| *ARR9-Q-R* | RT-qPCR | TGCTCTATCAGTTGAAATCCC |
| *ARR15-Q-F* | RT-qPCR | ATCTCCATCATCATCATCAAC |
| *ARR15-Q-R* | RT-qPCR | GACTCTAATTTGATCCTCTTGG |
| *ARR16-Q-F* | RT-qPCR | ATCATCATCTTCATCATGGTC |
| *ARR16-Q-R* | RT-qPCR | CAAGAGATCTTGAGCAACC |
| *CKX1-Q-F* | RT-qPCR | CTGAGAAGCGGAATTCTGAAC |
| *CKX1-Q-R* | RT-qPCR | GAGTACCCTGATCCATTTAACCA |
| *CKX2-Q-F* | RT-qPCR | TCCAACAAACCGGAATAAATG |
| *CKX2-Q-R* | RT-qPCR | TTGGGGTAGCGGATTGTAGT |
| *CKX3-Q-F* | RT-qPCR | TCTCAATACACAGTCAACGAGGA |
| *CKX3-Q-R* | RT-qPCR | TCGTACATAAACCCTCTTACATGG |
| *CKX4-Q-F* | RT-qPCR | GACACGTTAAGTAGAACTCTAG |
| *CKX4-Q-R* | RT-qPCR | CTTCCCATAGTCCTAAAGATC |
| *CKX5-Q-F* | RT-qPCR | CCATGGTCCTCAAATTAGTAACG |
| *CKX5-Q-R* | RT-qPCR | TCTGAGCATCTCATCACCTCTC |
| *CKX6-Q-F* | RT-qPCR | CAACTGGAGATTGTCACAGGAA |
| *CKX6-Q-R* | RT-qPCR | CCTAAACCACCAAGAACACCA |
| *CKX7-Q-F* | RT-qPCR | CACCAGAGCTAGGGTTTTGC |
| *CKX7-Q-R* | RT-qPCR | CATCGAACTCGGTGTATACTACTCTT |
| *CYP735A1-Q-F* | RT-qPCR | GGCCATGGTTTCGCAATC |
| *CYP735A1-Q-R* | RT-qPCR | CCGTTTCCGTTAAGCAAAGC |
| *CYP735A2-Q-F* | RT-qPCR | ATGGTGTCCCTTCCGTTGAACA |
| *CYP735A2-Q-R* | RT-qPCR | GAGGGTAAAGTCTTAATGACTCG |
| *NRT2.1-Q-F* | RT-qPCR | AACAAGGGCTAACGTGGATG |
| *NRT2.1-Q-R* | RT-qPCR | CTGTGGAAGGAGGCAAGAAC |
| *NRT2.4-Q-F* | RT-qPCR | GCTTGTGGAGCTACCTTTGC |
| *NRT2.4-Q-R* | RT-qPCR | GAAGTTTCCACCAGCTCCAG |
